# Do Arabidopsis *Squamosa promoter binding Protein-Like* genes act together in plant acclimation to copper or zinc deficiency?

**DOI:** 10.1101/590182

**Authors:** Anna Schulten, Lucas Bytomski, Julia Quintana, María Bernal, Ute Krämer

## Abstract

The genome of *Arabidopsis thaliana* encodes approximately 260 copper (Cu)-dependent proteins, which include enzymes in central pathways of photosynthesis, respiration and responses to environmental stress. Under Cu-deficient growth conditions, Squamosa promoter binding Protein-Like 7 (SPL7) activates the transcription of genes encoding Cu acquisition systems and mediates a metabolic reorganization to economize on Cu. The transcription factor SPL7 groups among comparably large proteins in the SPL family, which additionally comprises a second group of small SPL proteins targeted by miRNA156 with roles in plant development. SPL7 shares extended regions of sequence homology with SPL1 and SPL12. Therefore, we investigated the possibility of a functional overlap between these three members of the group of large SPL family proteins. We compared the *spl1 spl12* double mutant and the *spl1 spl7 spl12* triple mutant with the wild type and the *spl7* single mutant under normal and Cu-deficient growth conditions. Biomass production, chlorophyll content and tissue elemental composition at the seedling stage, as well as plant and flower morphology during reproductive stages, confirmed the involvement of SPL7, but provided no indication for important roles of SPL1 or SPL12, in the acclimation of Arabidopsis to Cu deficiency. Furthermore, we analyzed the effects of zinc (Zn) deficiency on the same set of mutants. Different from what is known in the green alga *Chlamydomonas reinhardtii*, Arabidopsis did not activate Cu deficiency responses under Zn deficiency, and there was no Cu overaccumulation in either shoot or root tissues of Zn-deficient wild-type plants. Known Zn deficiency responses were unaltered in *spl7, spl1 spl12* and *spl1 spl7 spl12* mutants. We observed that CuZnSOD activity is strongly downregulated in Zn-deficient *A. thaliana*, in association with an about 94% reduction in the abundance of the *CDS2* transcript, a known target of miR398. However, different from the known Cu deficiency responses of Arabidopsis, this Zn deficiency response was independent of *SPL7*and not associated with an upregulation of *MIR398b* primary transcript levels. Our data suggest that there is no conservation in *A. thaliana* of the crosstalk between Zn and Cu homeostasis mediated by the single SPL family protein CRR1 in Chlamydomonas. In the future, resolving how the specificity of SPL protein activation and recognition of target gene promoters is achieved will advance our understanding of the specific functions of different SPL family proteins in either Cu deficiency response regulation or growth and development of land plants.

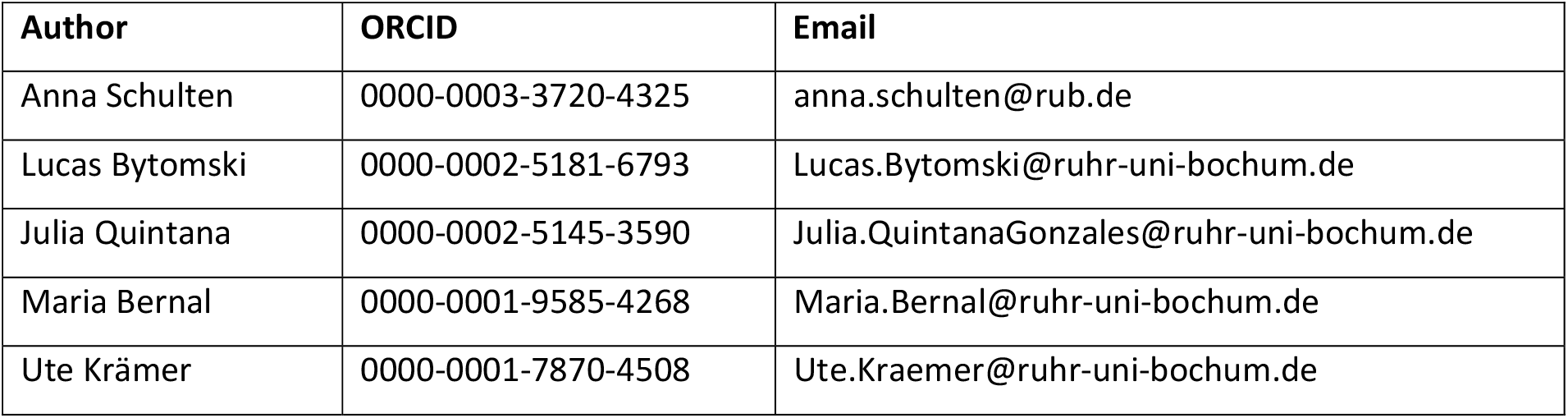

## 1 Introduction

Micronutrient metals act as cofactors in a multitude of proteins and are thus essential for plant growth, development and reproduction. Substantial proportions of the cellular quota of the micronutrient metals iron (Fe), zinc (Zn), manganese (Mn) and copper (Cu), for example, are allocated to proteins acting in photosynthesis (Yruela 2013), which places these metals at the core of plant energy metabolism and highlights their importance for plant-specific biochemistry. As sessile organisms, land plants depend entirely on their rhizosphere soils for the supply of nutrients, and they often experience nutrient imbalances. Not only are micronutrient metals essential but they can cause toxicity when present in excess. Thus, plants possess a molecular metal homeostasis network to adjust metal acquisition, distribution, utilization and storage to both the environmental conditions and the plant’s needs (Burkhead et al. 2009; Jeong and Guerinot 2009; Marschner and Marschner 2012; Sinclair and Krämer 2012).

A fundamental component of metal deficiency responses is the increased mobilization and uptake of scarce nutrients. For instance, Fe deficiency leads to the transcriptional upregulation of *Iron-Regulated Transporter 1 (IRT1)*, which encodes a transmembrane transporter mediating the uptake of Fe^2+^ from the soil solution into root cells (Eide et al. 1996; Vert et al. 2002). Since IRT1 is not specific for the transport of Fe^2+^ alone, other divalent metal cations such as Zn^2+^, Mn^2+^ and cobalt (Co^2+^) accumulate in root tissues of Fe-deficient plants (Vert et al. 2002; Cohen et al. 1998). Crosstalk between transition metals was also observed in Cu-deficient plants, which exhibited secondary physiological Fe deficiency because of a Cu-dependent defect in root-to-shoot Fe translocation (Bernal et al. 2012). A striking connection between Cu and Zn homeostasis was discovered in the green alga *Chlamydomonas reinhardtii*, in which Cu is overaccumulated under Zn deficiency (Hong-Hermesdorf et al. 2014). This is associated with the activation of Cu deficiency responses despite a high cellular Cu content. The Cu overaccumulation is dependent on the transcription factor Copper Response Regulator 1 (CRR1), which acts a master regulator of gene expression in response to Cu deficiency (Kropat et al. 2005; Hong-Hermesdorf et al. 2014).

In Arabidopsis, the closest homolog of CRR1 is Squamosa promoter binding Protein-Like 7 (SPL7) and is, similarly, the major known transcription factor mediating Cu deficiency responses (Yamasaki et al. 2009; Bernal et al. 2012). Both CRR1 and SPL7 activate the transcription of genes acting to increase cellular Cu uptake under Cu-deficient growth conditions. Additionally, CRR1 mediates the transcriptional upregulation of heme-containing Cytochrome *c* 6 (CYC6) to replace Cu-containing plastocyanin (PC) in the photosynthetic electron transport chain (Eriksson et al. 2004; Quinn and Merchant 1995). PC protein is concurrently degraded by a protease that is also under transcriptional control of CRR1 (Castruita et al. 2011). By contrast, PC is essential in *Arabidopsis thaliana* (Weigel et al. 2003). Instead, SPL7 activity results in the replacement of the abundant CuZn Superoxide Dismutases (CSD) with Fe Superoxide Dismutase 1 (FSD1) to economize on Cu (Yamasaki et al. 2007; Abdel-Ghany and Pilon 2008). Both SPL7 and CRR1 are members of the family of proteins characterized by the Squamosa promoter Binding Protein (SBP) domain, which consists of 76 highly conserved amino acids and contains both the nuclear localization signal and the recognition domain for the binding of a GTAC core DNA motif (Cardon et al. 1999; Birkenbihl et al. 2005; Yamasaki et al. 2006). So far, members of this protein family were exclusively found in the green plant lineage (Klein et al. 1996; Birkenbihl et al. 2005).

In *A. thaliana*, there are 16 SBP-Like (SPL) proteins, which group in two subfamilies based on size and sequence similarity (Guo et al. 2008; Xing et al. 2010). SPL1, SPL7, SPL12, SPL14 and SPL16 are more than 800 amino acids in length and are considered members of the group of large SPLs, whereas the remaining 11 SPLs are less than 400 amino acids long and are consequently addressed as small SPLs (Xing et al. 2010). Except for *SPL8*, small *SPLs* are targeted post-transcriptionally by miR156 (Schwab et al. 2005; Rhoades et al. 2002). The levels of miR156 decline with progressive plant development (Wang et al. 2009; Wu et al. 2009; Wu and Poethig 2006), which leads to a gradual increase in the expression of miR156-targeted SPLs. This constitutes a regulatory module in diverse developmental processes such as developmental phase transition, the specification of floral meristem identity, shoot branching and lateral root development (Wu and Poethig 2006; Schwarz et al. 2008; Wu et al. 2009; Yamaguchi et al. 2009; Yu et al. 2015; Xu et al. 2016; He et al. 2018).

Whereas *SPL7* is functionally well characterized (Yamasaki et al. 2009; Bernal et al. 2012), much less is known about the biological roles of other large SPLs. Arabidopsis *spl14* mutants showed increased sensitivity to the fungal toxin fumonisin B1 and altered plant architecture, namely elongated petioles and enhanced serration of the leaf margins (Stone et al. 2005). SPL1 and SPL12 were recently implicated in thermotolerance, especially of reproductive tissues (Chao et al. 2017). Whereas proteins of the SPL family generally show little amino acid sequence conservation outside of the SBP domain, SPL1 and SPL12 share additional regions of sequence homology with SPL7. These comprise a region C-terminal of the SBP domain containing the sequence WL(X)_3_P(X)_3_E(X)_2_IRPGC that is also conserved in CRR1 (Cardon et al. 1999; Döring et al. 2000; Kropat et al. 2005). In addition, SPL7, SPL1 and SPL12 all contain an AHA motif at the N-terminus, which is thought to act as a transcriptional activator domain and is also present in the long SPLs SPL14 and SPL16. Although there has been continuous progress in understanding the distinct biological roles of individual SPL proteins (Xu et al. 2016), we still know only little about how target gene specificity is achieved, especially considering the highly conserved DNA-binding domain across the entire SPL protein family.

Because functional overlap and the ability to compensate for the loss of other *SPL* gene functions have been reported for several SPL family members (Wu and Poethig 2006; Xing et al. 2010; Xing et al. 2013; Xu et al. 2016), it is conceivable that additional *SPL* genes may have partially overlapping biological functions. We hypothesized that *SPL1* and *SPL12* might have functions related to those of *SPL7* or *CRR1* in the regulation of transition metal homeostasis. In response to acute heat stress (1h at 42 °C), the putative Cu exporter-encoding gene *HMA5* is transcriptionally upregulated in the wild type but not in an *spl1 spl12* double mutant (Andrès-Colàs et al. 2006; Burkhead et al. 2009; Chao et al. 2017). This observation provided circumstantial support for a possible association of *SPL1* and *SPL12* with Cu homeostasis.

In this study, we addressed two questions. First, is there any evidence for functions of *SPL1* and *SPL12* in the responses of Arabidopsis to Cu or Zn deficiency? Second, does Zn deficiency result in Cu over-accumulation in *A. thaliana*, and is this dependent on *SPL7*, as known in *C. reinhardtii?*

## 2 Methods

### Plant material

*Arabidopsis thaliana* wild-type seeds (Col-0) were obtained from Lehle seeds. The double mutant *spl1 spl12* and the triple mutant *spl1 spl7 spl12* were kindly provided by Dr. Peter Huijser (Max Planck Institute for Plant Breeding Research, Cologne, Germany) and are crosses of the T-DNA insertion lines SALK_070086 *(spl1)*, SALK_017778 *(spl12)* and SALK_125385 (spl7-2), obtained from the Nottingham Arabidopsis Stock Centre. The *spl7-2* mutant was characterized earlier (Bernal et al. 2012). Plants homozygous for the T-DNA insertions were identified by PCR. For detection of the T-DNA insertion alleles, the primers At_spl1geno_f and At_spl12geno_f were used in combination with the primer specific for the left border of the T-DNA (see Supplemental Table 1), respectively.

### Growth conditions

Most laboratories generate micronutrient metal deficiencies by including an excess of a chelator in order to render contaminant metals unavailable for plants during the cultivation period. Instead, we generate metal deficiencies by removing contaminant metals from the media so that plants can be cultivated in the absence of a chelator excess (Quinn and Merchant 1998; Salomé et al. 2014). For experiments in sterile culture on petri dishes, wild-type or mutant seeds were surface sterilized and sown on modified Hoagland’s medium (Becher et al. 2004) containing 1% (w/v) sucrose and solidified with 1% (w/v) EDTA-washed Agar Type M (Sigma-Aldrich, Steinheim, Germany), followed by stratification in the dark at 4 °C for 2 d. EDTA-washed agar was prepared by stirring the required amount of agar including a 15% surplus in 10 mM EDTA pH 5.7 for 1 d. After another two identical washes in EDTA solution, the agar was washed once with ultrapure water for 1 d and five times with ultrapure water for 1 h each in order to remove all remaining EDTA. The volume of the wash solution for each wash step corresponded to the desired final volume of medium. Wash solutions were removed by decanting after the agar had sedimented by gravity in between each step. To account for remaining water in the wet agar after the washes, half of the final volume 2X modified Hoagland’s medium was prepared, mixed with the wet agar and then filled up to final volume (1X) with ultrapure water.

20 seedlings were cultivated on each vertically orientated round glass petri dish (diameter of 150 mm) for Cu deficiency experiments or on each square polypropylene petri dish (120 mm × 120 mm) for Zn deficiency experiments in an 11 h day (145 μmol m^−2^ s^−1^, 22 °C)/ 13 h night (18 °C) cycle in a growth chamber (CLF Plant Climatics, Wertingen, Germany) for 21 d. Glass petri dishes were soaked in 0.2 N HCl overnight and rinsed with deionized water before autoclaving. The positions of petri dishes within the growth chamber were randomized once per week. For soil cultivation, plants were grown in commercially available standard soil Type Minitray (Balster Einheitserdewerk, Fröndenberg, Germany). After stratification in darkness at 4 °C for 2 d, plants were transferred to a growth chamber (CLF Plant Climatics, Wertingen, Germany) and grown in a 16 h day (145 μmol m^−2^ s^−1^, 22 °C), 8 h night (18 °C) cycle. For each experiment, half of the plants were watered with 2 mM CuSO_4_ in tap water (+Cu) once per week, whereas the other half received tap water without additional CuSO_4_ (-Cu). The positions of plant pots within trays and tray positions within the growth chamber were randomized once per week.

### Determination of plant biomass and elemental concentrations

Shoots and roots of 21-day-old plate-grown seedlings were separated with a scalpel, pooled from 20 seedlings, then washed in ultrapure water and carefully blotted dry to determine fresh biomass. Subsequently, extracellularly bound metal cations were desorbed from pooled shoot and root tissues as follows: 10 min in 2 mM CaSO_4_, 10 mM EDTA, 1 mM MES pH 5.7; 3 min in 0.3 mM bathophenanthroline disulphonate, 5.7 mM sodium dithionite, 1 mM MES pH 5.7; twice for 1 min in ultrapure water (Cailliatte et al. 2010, with modifications). Shoot and root tissues were then dried at 60 °C for 3 d and equilibrated at RT for at least 3 d before homogenization. The digestion of plant material and multi-element analysis of digests was performed as described (Sinclair et al. 2017).

### Measurement of chlorophyll concentrations

Total chlorophyll was extracted with 2 mL methanol from 20 mg of frozen ground shoot material of 21-day-old seedlings. Extinction values were determined spectrophotometrically in 96-well plates and path length-corrected to 1 cm with the factor 0.51 (Warren 2008), and chlorophyll concentrations calculated as described (Porra et al. 1989).

### RNA extraction and quantitative real time RT-PCR

For RNA extraction, all harvests were performed at ZT = 3.5 (3.5 h after lights on). Shoot tissues were pooled of 20 seedlings per plate, immediately shock-frozen in liquid nitrogen and homogenized by grinding. RNA was isolated from 50 mg-subsamples of shoot tissue using TRIzol reagent (Thermo Fisher, Schwerte, Germany) according to manufacturer’s instructions. Equal amounts of RNA were treated with DNase I with the TURBO DNA-free Kit (Thermo Fisher, Schwerte, Germany). One μg RNA was used for cDNA synthesis with oligo-dT primers and the Revert Aid First Strand synthesis kit (Thermo Fisher, Schwerte, Germany). Four μL of diluted cDNA (equivalent to 8 ng of RNA per well) were used as template for quantitative real time RT-PCRs which were performed in 384-well plates using a LightCycler 480 II detection system (Roche Diagnostics, Mannheim, Germany). GoTaq PCR Mastermix (Promega, Mannheim, Germany) was used to monitor cDNA amplification. Reaction efficiencies (RE) and C_T_ values for each PCR reaction were determined with the LinRegPCR program, version 2016.0 (Ruijter et al. 2009; Tuomi et al. 2010; Ruijter et al. 2014). Relative transcript levels (RTL, in ‰ of the reference gene) were calculated as follows: RTL = 1000 × RE_m_^−ΔCT^, with RE_m_ as the mean of reaction efficiencies per primer pair and ΔC_T_ = C_T_ (target gene) – C_T_ (constitutively expressed reference gene: *EF1α*), as described (Bernal et al. 2012). Primer sequences are listed in Supplemental Table 1.

### In-gel detection of superoxide dismutase activities

Shoot tissues were homogenized in liquid nitrogen. Soluble native proteins were extracted from 100 mg fresh biomass of ground shoot material by vortexing in extraction buffer (50 mM Tris-HCl pH 7.4, 1% (w/v) PVP-40, 1 mM EDTA, 1 mM PMSF, 5 mM DTT). Total protein was quantified in the supernatant using the Bradford assay (Bradford 1976), employing bovine serum albumin as a standard for calibration. Superoxide dismutase (SOD) activity was determined semi-quantitatively using an in-gel staining method (Beauchamp and Fridovich 1971). Twenty μg of total protein per sample were separated on native 12% (v/v) polyacrylamide gels. The staining was essentially done as described (Beauchamp and Fridovich 1971), modified by using 1.2 mM NBT and 21 mM TEMED and by performing two additional gel washes with potassium phosphate buffer (36 mM, pH 7.8) before light exposure. Specific inhibitors were used during the activity staining period in replicate gels (data not shown) for the identification of different SOD isoforms (3 mM H_2_O_2_ to inhibit both FeSOD and CuZnSOD, 0.1 mM KCN to inhibit CuZnSOD activities (Sandalio et al. 1987)).

### Accession numbers

Sequence data for the genes mentioned in this article can be found on The Arabidopsis Genome Initiative (TAIR) or GenBank website. The AGI locus identifiers are listed in Supplemental Table 2.

## 3 Results

In order to address the possibility of a functional overlap between paralogous members of the SPL protein family, we compared Cu and Zn deficiency responses of the *spl1 spl12* double and the *spl1 spl7 spl12* triple mutant to the wild type (WT) and the *spl7* single mutant. We confirmed by RT-PCR that *SPL1* and *SPL12* were expressed in 21-day old WT seedlings under both Cu- and Zn-deficient growth conditions (Fig. S1), which we generated by removing trace metal contaminations arising from agar and glassware and by omitting Cu and Zn, respectively, from the synthetic growth media (Schulten and Krämer 2018). In addition, we confirmed that full-length *SPL1* and *SPL12* transcripts were absent or reduced to nearly undetectable levels under all our growth conditions in the mutants *spl1 spl12* and *spl1 spl7 spl12* (Fig. S1).

### 3.1 Role of *SPL1* and *SPL12* in the Cu deficiency response at seedling stage

The previously described growth defects of the *spl7* single mutant under Cu deficiency were evident through strongly reduced fresh biomass and lower chlorophyll concentrations (Fig. 1A-C, Fig. S2). Moreover, the *spl1 spl7 spl12* triple mutant mimicked *spl7* whereas the *spl1 spl12* mutant performed like the wild type in media lacking added Cu (Fig. 1A-C, Fig. S2). The *spl1 spl7 spl12* mutant only differed from *spl7* through lower chlorophyll concentrations under-Cu. As expected, Cu concentrations around or below 2 μg g^−1^ shoot DW confirmed deficiency in all plants cultivated in −Cu media (Fig. 1D). Under Cu-sufficient growth conditions, Cu concentrations were clearly reduced in rosettes of both *spl7* and *spl1 spl7 spl12* (around 4 μg g^−1^ DW) compared to 8 μg Cu g^−1^ shoot DW in the wild type (Fig. 1D). It might seem surprising that shoot Cu concentrations of seedlings cultivated under Cu-deficient conditions were similar in WT, *spl7* and *spl1 spl7 spl12* (Fig. 1D). However, biomass production is Cu-limited in plants that cannot activate SPL7-dependent Cu deficiency responses (see Fig. 1B), thus counteracting an additional decrease in tissue Cu concentrations (Bernal et al. 2012). Our observation that there is no growth impairment in wild-type and *spl1 spl12* mutant seedlings containing as little as 1 μg Cu g^−1^ DW may result from Cu economization as part of *SPL7* functions impacting subcellular Cu distribution (Abdel-Ghany et al. 2005; Abdel-Ghany 2009; Bernal et al. 2012). To further investigate the functionality of the Cu economy response, we performed an in-gel staining for Superoxide Dismutase (SOD) activities. When cultivated in Cu-deficient medium, the wild type showed strongly enhanced FeSOD (FSD) activity and dramatically reduced CuZnSOD (CSD) activity, as expected (Bernal et al. 2012). The activity pattern was nearly identical to the wild type in *spl1 spl12* under both +Cu and −Cu (Fig. 1E). By contrast, when grown in Cu-deficient medium, both *spl7* and *spl1 spl7 spl12* lacked FSD activity and activated Mn Superoxide Dismutase 1 (MSD1) instead, in addition to aberrant, clearly detectable levels of residual CSD activity. By comparison, the altered SOD activity patterns in *spl1 spl7 spl12* seemed to result solely from the loss of *SPL7* function. Unlike FSD and CSD, MSD1 activity has not been directly linked to the transcription factor activity of SPL7 so far and might be increased here as a result of a secondary response to maintain superoxide scavenging. In summary, these data provided no indication for an important role of *SPL1* or *SPL12* in Cu deficiency responses at the seedling stage.

**Figure 1:**
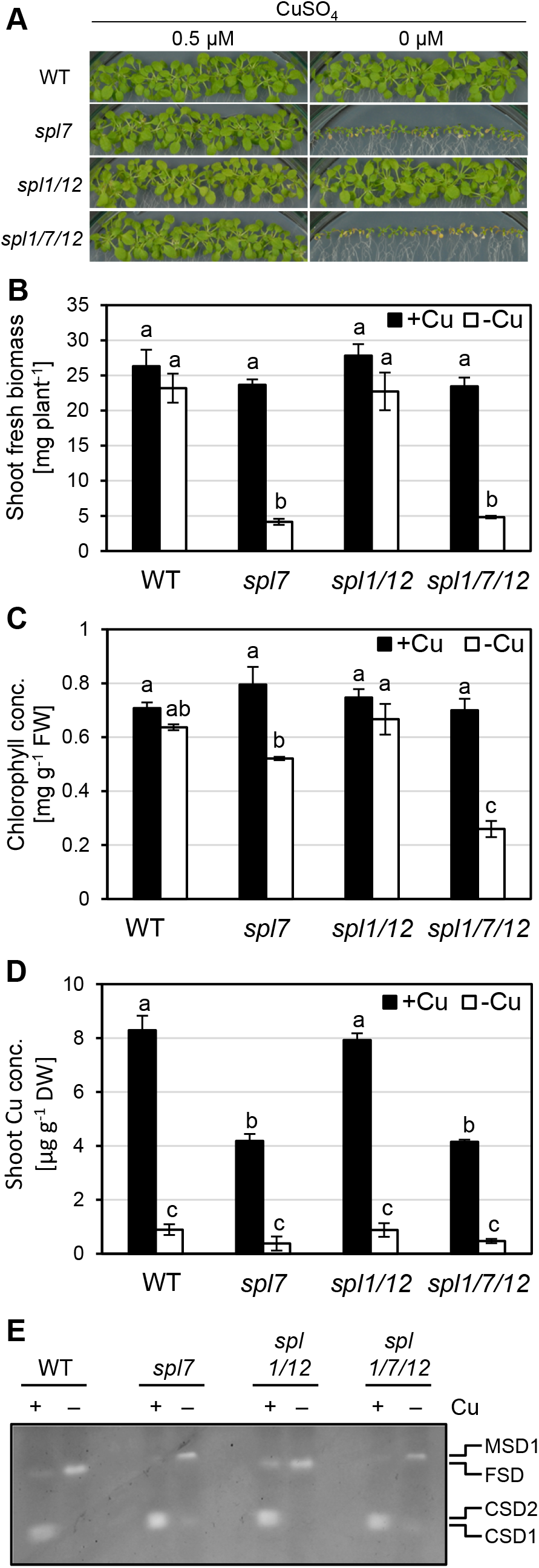
Comparison of copper deficiency symptoms in wild-type, *spl7* single, *spll spl12* double and *spll spl7 spl12* triple mutant Arabidopsis seedlings. (A) Photographs of 21-day-old seedlings grown on Cu-sufficient (0.5 μM CuSO_4_) or Cu-deficient (no added CuSO_4_) agar-solidified media in vertically oriented glass plates in short days (11 h). (B-D) Fresh biomass (B), chlorophyll concentration (C) and Cu concentration (D) of shoots of seedlings as described in (A). Bars represent arithmetic means ± SD (*n* = 3 replicate plates, each with 20 seedlings). Different characters denote statistically significant differences (*P* < 0.05) between means based on ANOVA (Tukey’s HSD). (E) In-gel detection of superoxide dismutase activities in total protein extracts of seedlings grown as described in (A). Total protein (20 μg per lane) was separated in a 12% (v/v) native polyacrylamide gel. CSD1, CSD2: Cu/Zn Superoxide Dismutases, FSD: Fe Superoxide Dismutase 1-3, MSD1: Mn Superoxide Dismutase 1. Data are from one experiment representative of two to three independent experiments.

### 3.2 Role of *SPL1* and *SPL12* in the Cu deficiency response at reproductive stage

Since the involvement of *SPL1* and *SPL12* in thermotolerance was observed at the reproductive stage and specifically in inflorescences (Chao et al. 2017), we next monitored plant and flower morphology of *spl1 spl12* and *spl1 spl7 spl12* in dependence on Cu supply at this later developmental stage upon soil cultivation (Fig. 2). On our unsupplemented soil, we observed stunted growth and withered flowers in *spl7* (Fig. 2G-H, Fig. 3A). Therefore, we suspected that our soil is low in bioavailable Cu (-Cu), and we generated an additional Cu-sufficient growth condition by watering with 2 mM CuSO_4_ (+Cu). With this treatment, the *spl7* phenotype can be partially rescued (Fig. 2E-F, Fig. 3), in agreement with a significantly higher rosette Cu concentration compared to the −Cu condition (Fig. S3A). Compared to the wild type, rosette Zn concentrations were generally slightly decreased by about 30% to ca. 35 μg g^−1^ DW in *spl7* and the triple mutant. This effect may be unrelated to *spl7*, however, because Zn concentrations were independent of Cu supply in all genotypes (Fig. S3B), whereas SPL7 transcription factor activity is thought to be strongly enhanced under Cu deficiency.

**Figure 2:**
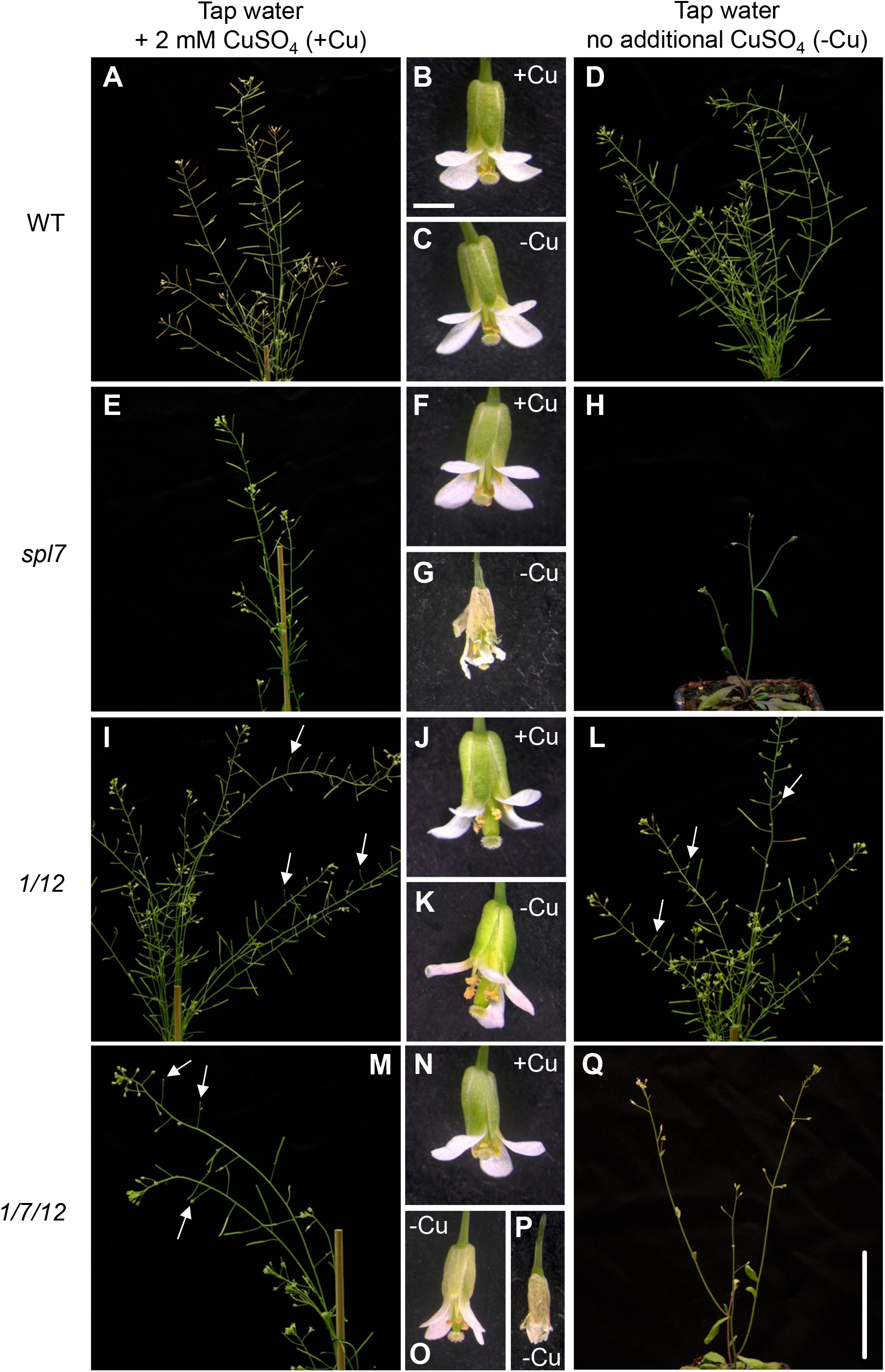
Photographs of reproductive-stage wild-type, *spl7* single, *spl1 spl12* double and *spl1 spl7 spl12* triple mutant Arabidopsis plants cultivated on Cu-deficient and -sufficient soil. (A-H) Morphology of 40-day-old plants and their flowers. Plants were watered with equal amounts of either tap water or 2 mM CuSO_4_ in tap water (freshly prepared) and grown in long days (16 h). Photographs are from one experiment (6 replicate plants per genotype and treatment) representative of two independent experiments. Scale bars (Q) correspond to 5 cm (A, D, E, H, I, L, M, Q), or (B) to 1 mm (B, C, F, G, J, K, N, O). White arrows highlight the positions of aborted siliques.

Plant morphology at the reproductive stage was unaffected by Cu supply in both the wild type and *spl1 spl12* (Fig. 2A-D, I-L). The *spl1 spl12* mutant displayed partial sterility. Some flowers along the inflorescences did not develop into siliques (Fig. 2I, L), and this was also reflected in a lower number of siliques per plant and seeds per silique compared to the WT (Fig. 3B, C). This phenotype was previously reported (Chao et al. 2017), and it was independent of Cu supply under our growth conditions (Fig. 2I-L, Fig. 3). Under +Cu conditions, the overall morphology of the *spl1 spl7 spl12* triple mutant was intermediate and combined aspects of *spl7* and *spl1 spl12* (Fig. 2E, M). Under −Cu, the triple mutant was similar to *spl7* with strongly reduced plant size and reduced branching of the inflorescence, yet it grew slightly larger than *spl7* (Fig. 2H, Q). Notably, *spl1 spl7 spl12* grown under −Cu had withered flowers on the main inflorescence stem but healthy flowers on the lateral stems (Fig. 2O-Q). This is unlike *spl7* cultivated in Cu-deficient soil, which solely produced withered, unhealthy flowers (Fig. 2G-H) and only rarely developed any siliques at all (Fig. 3A, B). The proportion of plants capable of producing siliques on −Cu soil was about four-fold higher for *spl1 spl7 spl12* than for *spl7*, but it was not fully restored to the levels in WT or *spl1 spl12* (Fig. 3A). The number of siliques developed per plant was similarly low in the triple mutant as in *spl7* (Fig. 3B). The number of seeds per silique was even lower in the triple mutant than in *spl7*, raising the possibility that *SPL1/SPL12* and *SPL7* have additive effects on seed formation under +Cu growth conditions, possibly through differing mechanisms (Fig. 3B-C). Under −Cu conditions, the difference was even more pronounced, consistent with a small contribution of *SPL1/SPL12* to Cu deficiency acclimation in the absence of *SPL7* only. Taken together, these results suggest overall complex interactions of *spl7* with *spl1* and *spl12*, which have comparably minor phenotypic effects. The largest of these effects is an apparent antagonism between *SPL7* and the pair of *SPL1* and *SPL12*, or one of these two genes. In summary, this experiment provided no unequivocal evidence for a role of *SPL1* or *SPL12* similar to *SPL7* in alleviating Cu deficiency symptoms during the reproductive phase of development.

**Figure 3:**
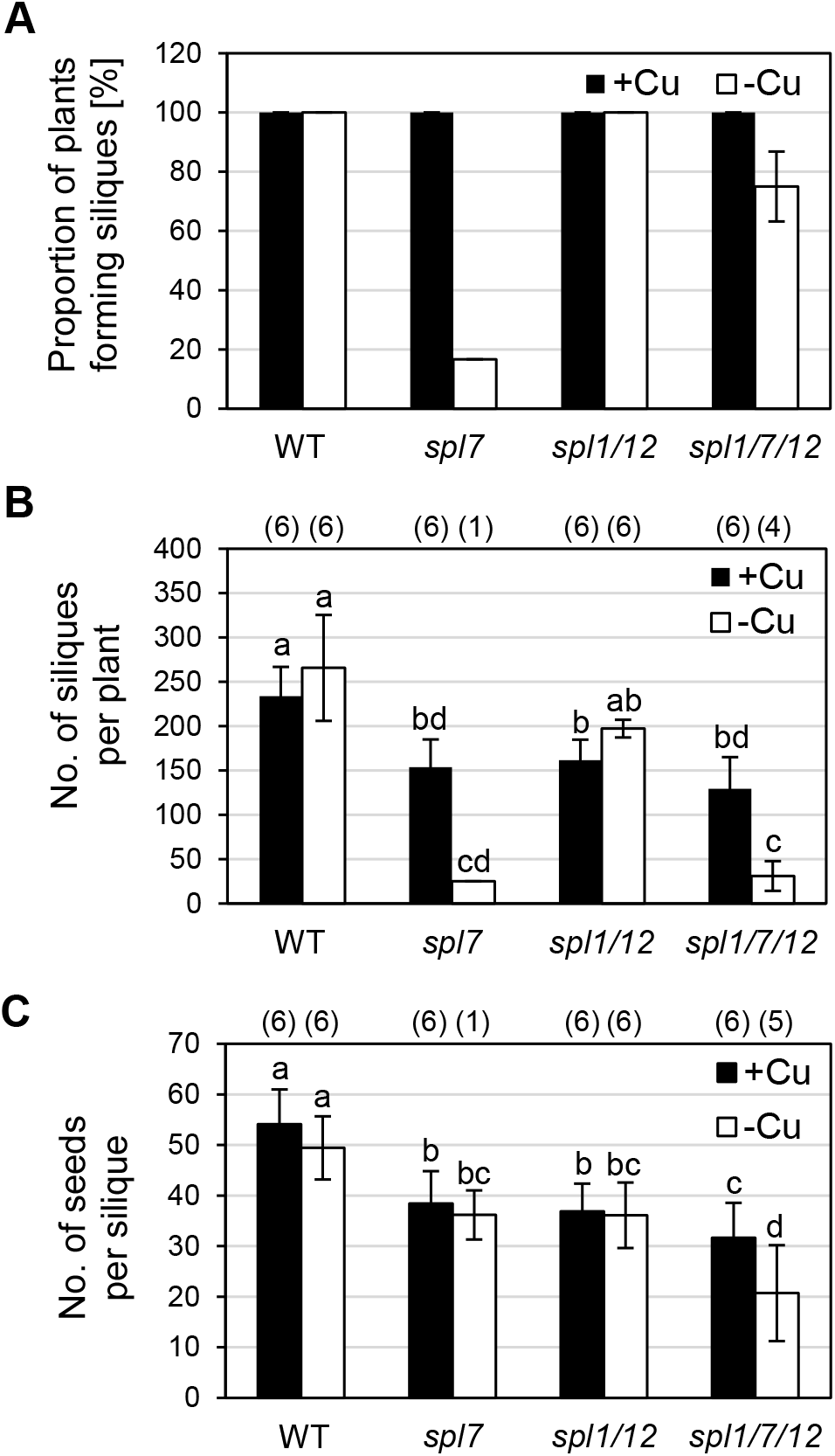
Seed production in wild-type, *spl7* single, *spl1 spl12* double and *spl1 spl7 spl12* triple mutant Arabidopsis plants cultivated on Cu-deficient and -sufficient soil. (A) Proportion of plants that produced siliques (length > 10 mm). Bars represent arithmetic means ± SD (n = 2 percentage values, each from one independent experiment with six replicate plants per genotype and treatment). (B) Number of siliques (length > 10 mm) per plant. Bars show arithmetic means ± SD (n = 1 to 6 plants that produced siliques per genotype and treatment in one experiment representative of two independent biological experiments; see numbers in parentheses above the bars and (A) for the number of replicate plants). (C) Number of seeds per silique. Bars represent arithmetic means ± SD (n = 6 to 36, i.e. 6 siliques for each plant that produced siliques [length > 10 mm] out of 6 replicate plants; see numbers in parentheses above the bars and (A) for the number of replicate plants) from one experiment representative of two independent experiments. Different characters denote statistically significant differences (P < 0.05) between means based on ANOVA (Tukey’s HSD). Plants were grown as described in Fig. 2, and siliques were counted on 40-day-old plants.

### 3.3 Role of *SPL1, SPL7* and *SPL12* in the Zn deficiency response

Under Zn deficiency, Chlamydomonas was reported to exhibit CRR1-dependent Cu overaccumulation alongside physiological Cu deficiency, as indicated by the transcriptional upregulation of the CRR1 target *CYC6* as well as a decreased abundance of plastocyanin protein (Hong-Hermesdorf et al. 2014). Therefore, we examined the effect of Zn deficiency on Cu homeostasis in wild-type Arabidopsis and mutants in *CRR1* homologues. Under Zn-deficient growth conditions in sterile culture, WT, *spl7, spl1 spl12* and *spl1 spl7 spl12* mutant seedlings all showed similarly reduced shoot and root fresh biomass, similarly reduced chlorophyll content and strongly and similarly decreased shoot and root Zn concentrations when compared to seedlings grown under Zn sufficiency (Fig. 4, Fig. S4). Based on root and shoot Cu concentrations, we observed only very minor increases (≤ 6%, not statistically significant) in Cu accumulation in roots or shoots of wild-type seedlings under Zn deficiency (Fig. 5A-B). As expected, Cu levels in both *spl7* and the triple mutant were lower than in the wild type, and the magnitude of this difference was independent of external Zn supply (Fig. 5A-B). We further examined metal homeostasis through in-gel staining for superoxide dismutase activities. Under Zn deficiency, no activity of CSD1 and CSD2 was detectable in any of the genotypes, and there was no compensatory upregulation of the activity of another SOD (Fig. 5C). Posttranscriptional downregulation of CSDs through miR398 is a characteristic of the SPL7-dependent Cu deficiency response (Yamasaki et al. 2009; Bernal et al. 2012). Therefore, we determined the transcript abundances of Cu and Zn deficiency markers in shoots of Zn-deficient seedlings (Fig. 6). Transcripts of the Zn deficiency markers *Zinc-Regulated Transporter*, *Iron-Regulated Transporter (ZRT-IRT)-like protein 9 (ZIP9)* and *Nicotianamine Synthase 2 (NAS2)* (Talke et al. 2006) were undetectable under Zn-sufficient conditions and similarly highly upregulated in all genotypes under Zn-deficient growth conditions (Fig. 6A), in accordance with severe Zn deficiency (see Fig. 4). Transcript levels of Cu deficiency markers *FSD1* and *MIR398b* were consistent with a more Cu-sufficient physiological status of Zn-deficient wild-type plants (Fig. 6B), in contrast to a Cu-deficient physiological status in Zn-deficient *C. reinhardtii* (Hong-Hermesdorf et al. 2014). The observation of a decrease in *miR398b* precursor transcript levels under Zn deficiency compared to control conditions in the wild type could be predicted to result in a concomitant increase in transcript levels of the *miR398* target *CSD2* under Zn deficiency. We observed the contrary: *CSD2* transcript levels were strongly decreased under Zn deficiency to 6% of those under Zn-deficient conditions in both the wild type and *spl7* (Fig. 6B), fully consistent with the profile of SOD activities (see Fig. 5C). This indicated that *CSD2* transcript levels are down-regulated under Zn-deficiency through a mechanism independent of *SPL7* and *miR398b*, in accordance with the requirement for Zn as a structural cofactor in CSD enzymes (Sawada et al. 1972; Richardson et al. 1975). In conclusion, our results suggest that the crosstalk between Zn and Cu deficiency reported for the green alga *C. reinhardtii* is not conserved in the land plant *A. thaliana.*

**Figure 4:**
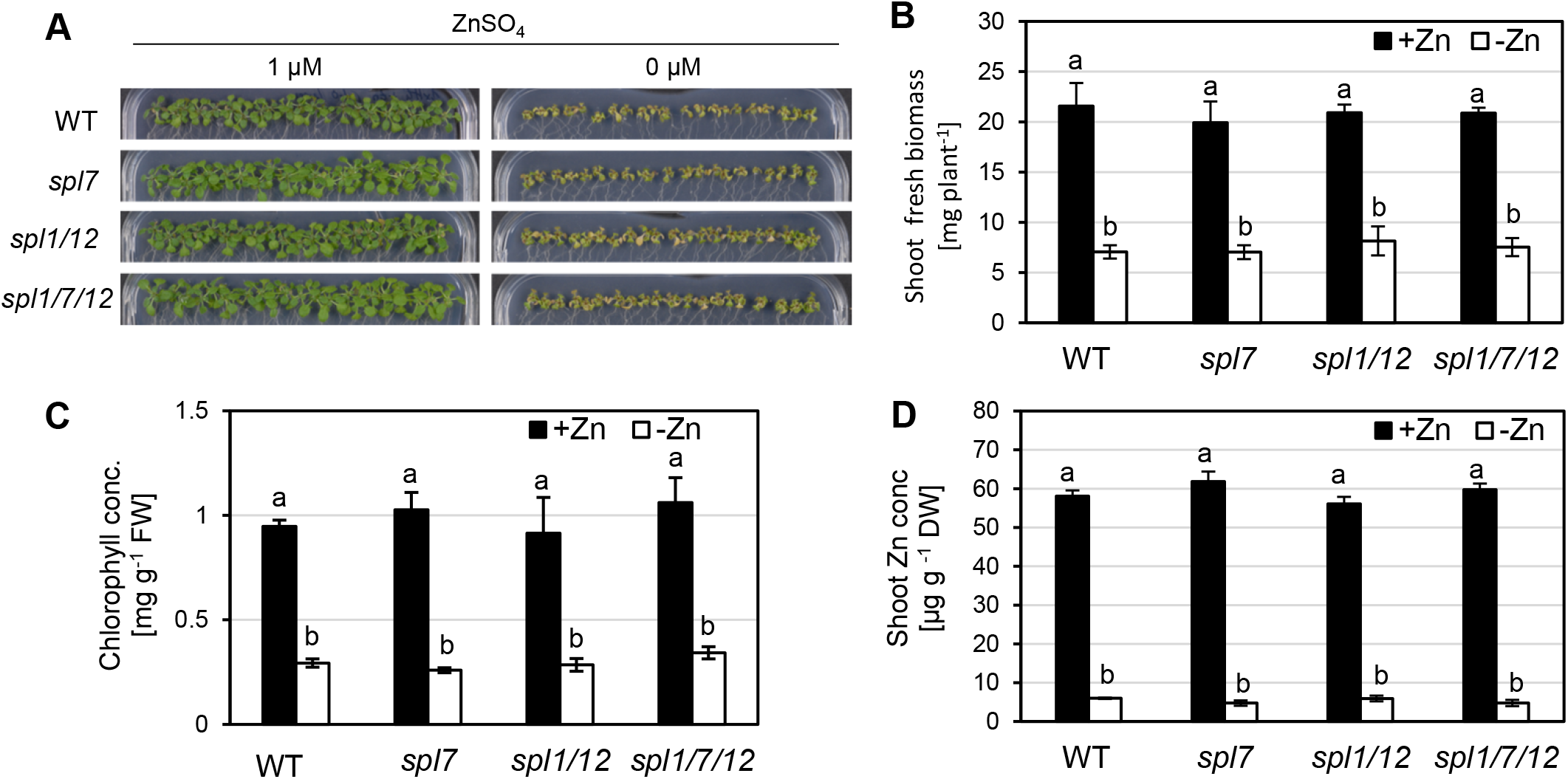
Comparison of zinc deficiency symptoms in wild-type, *spl7* single, *spl1 spl12* double and *spl1 spl7 spl12* triple mutant Arabidopsis seedlings. (A) Photographs of 21-day-old seedlings grown on Zn-sufficient (1 μM ZnSO_4_) or Zn-deficient (no added ZnSO_4_) agar-solidified media in vertically-oriented plastic petri dishes in short days (11 h). Photographs shown are from one experiment representative of three independent experiments. (B-D) Fresh biomass (B), chlorophyll concentration (C) and Zn concentration (D) for shoots of seedlings grown as described in (A). Bars represent arithmetic means ± SD *(n* = 3 replicate plates, each with 20 seedlings). Different characters denote statistically significant differences (P < 0.05) between means based on ANOVA (Tukey’s HSD). Data are from one experiment representative of two to three independent experiments.

## 4 Discussion

### 4.1 *SPL1* and *SPL12* are not involved in Cu deficiency responses

The role of *SPL7* in the Cu deficiency response of *Arabidopsis thaliana* is well established, but according to genome-wide transcriptomics only 13% of about 1,500 Cu deficiency-responsive genes were regulated in an SPL7-dependent manner (Yamasaki et al. 2009; Bernal et al. 2012; Garcia-Molina et al. 2014). This suggested that additional transcription factors contribute to Cu deficiency responses. The Cu deficiency-induced Transcription Factor 1 (CITF1) was identified to have roles in Cu uptake into roots and Cu delivery to flowers (Yan et al. 2017). *CITF1* transcript levels were increased under Cu deficiency, and this was entirely dependent on *SPL7* in leaves and partially dependent on *SPL7* in roots and flowers (Yan et al. 2017). While CITF1 activity might thus account for at least some of the identified SPL7-independent transcriptional responses to Cu deficiency, additional transcriptional regulators must be involved. Because of substantial functional redundancy observed among small SPL proteins (Xu et al. 2016), we hypothesized that some large SPL proteins might have SPL7-related functions under Cu deficiency. Given the presence of additional regions of sequence homology with SPL7 outside of the SBP domain (Cardon et al. 1999; Kropat et al. 2005), we considered SPL1 and SPL12 as large SPL protein candidates for such roles. SPL7 was proposed to have a dual role not only as a transcriptional activator of Cu deficiency responses but also as a Cu sensor protein (Sommer et al. 2010; Garcia-Molina et al. 2014). According to the amino acid sequence motifs present in the proteins, SPL1 and SPL12 could equally act in this manner. However, we observed no Cu deficiency-related phenotypes in the *spl1 spl12* mutant compared to WT. Moreover, *spl1 spl7 spl12* did not differ from *spl7* at seedling stage (Fig. 1). In comparison, a *citf1 spl7* double mutant was seedling lethal unless the mutant was fertilized extensively with Cu (Yan et al. 2017). Furthermore, the comparison of internal Cu concentrations, of the Cu economy response and of mutant phenotypes at the reproductive stage led us to conclude that *SPL1* and *SPL12* have no, or only very small, autonomous roles in plant acclimation to Cu deficiency, different from *SPL7* (Figs. 1–3, S1-S3). The *SPL7* transcript is detectable regardless of plant physiological Cu status, and consequently post-translational mechanisms were proposed to regulate SPL7 protein activity (Yamasaki et al. 2009; Garcia-Molina et al. 2014). Based on earlier *in vitro* studies, Cu+ ions can displace the Zn^2+^ ions bound to the Zn finger-like motifs of the SBP domain of CrCRR1 (Kropat et al. 2005; Sommer et al. 2010). The authors suggested a model in which an active, DNA-binding form of CRR1 containing Zn^2+^ is present under Cu deficiency *in vivo*, whereas under Cu sufficiency Cu itself acts as a signal that inactivates CRR1. In conflict with this model, the introduction of a transgene encoding a truncated SPL7 protein, which contained only the N-terminal part of SPL7 including the SBP domain, resulted in the constitutive transcriptional activation of SPL7 target genes irrespective of cellular Cu levels in the genetic background of the *spl7* mutant (Garcia-Molina et al. 2014). This observation is consistent with a role for the putative C-terminal transmembrane helix of SPL7 in the Cu-dependent regulation of SPL7 activity. It was proposed that the inactive protein is anchored to an endosomal membrane until, under Cu deficiency, an N-terminal moiety of SPL7 of approximately 45 kDa in size is released from the endosomal membrane via proteolytic cleavage by an unidentified protease (Garcia-Molina et al. 2014). Interestingly, all members of the group of large SPL proteins, including also SPL1 and SPL12, are predicted to comprise a C-terminal transmembrane helix by Aramemnon consensus prediction (Schwacke et al. 2003). However, the possibility of membrane tethering controlling transcription factor activity has not been proposed for any other large SPL proteins yet. SPL7 homodimerisation was also considered as a possible mechanism that regulates SPL7 activity (Garcia-Molina et al. 2014). The putative homodimerization domain identified in a yeast two-hybrid screen is conserved in both SPL1 and SPL12 (Cardon et al. 1999). All of the models recapitulated here might constitute general regulatory features common to all large SPL proteins rather than a specific characteristic of SPL7 or CrCRR1 and Cu deficiency responses. This highlights the importance of elucidating how target specificity is achieved among members of the SPL family in order to advance our understanding of their distinct functions and the mechanism of Cu sensing and how this is transduced to modulate the activity of SPL7.

Surprisingly, flower health and the proportion of plants capable of producing siliques were alleviated in the *spl1 spl7 spl12* triple mutant compared to *spl7* under Cu deficiency (Figs. 2 and 3). Possible antagonistic roles of these SPL proteins thus warrant further examination in the future. Moreover, it will be interesting to study the interplay of small and large SPLs in securing the fertility of both male and female floral organs, considering that the SPL8 and miR156-targeted small SPLs act to regulate anther development and gynoecium patterning, among other roles (Xing et al. 2010; Xing et al. 2013).

### 4.2 No role of *SPL1, SPL7* and *SPL12*, and no overaccumulation of Cu, in the acclimation of Arabidopsis plants to Zn deficiency

Different from the green alga Chlamydomonas, we found no evidence for a role of the *CrCRR1* homologues *SPL1, SPL7* or *SPL12* in Zn deficiency responses (Fig. 4). Moreover, in Chlamydomonas, Zn deficiency led to a dramatic overaccumulation of Cu. We did not observe such a response in *A. thaliana*, and Zn supply-dependent changes in Zn and Cu concentration in shoots and roots were unaltered in *spl7, spl1 spl12* or *spl1 spl7 spl12* (Figs. 5 and 6). In *C. reinhardtii*, the CRR1-dependent Cu accumulation was suggested to prevent the mis-metallation of Zn-requiring proteins under Zn deficiency through the sequestration of Cu in lysosome-related organelles called cuprosomes (Hong-Hermesdorf et al. 2014). Since the Cu stored in cuprosomes remains bioavailable, the possibility was raised that a selective advantage arises if nutrient deficiencies are encountered frequently. As a unicellular organism, Chlamydomonas might have to rely on strategies for metal distribution and storage different from vascular plants, which can distribute metals not only sub-cellularly but also across tissues and organs under conditions of nutrient imbalances. Indeed, the Cu-transporting Heavy Metal P-type ATPase HMA5 was hypothesized to have a role in basal Cu tolerance by exporting Cu from the root symplast into the apoplastic xylem for root-to-shoot transport (Andrès-Colàs et al. 2006; Kobayashi et al. 2008). Several members of the family of Yellow Stripe-Like (YSL) transporters were also implicated in intra-plant partitioning of metals including Cu, likely through the transport across membranes of metal nicotianamine chelates in land plants (DiDonato et al. 2004; Curie et al. 2009; Zheng et al. 2012). Both the YSL protein family and nicotianamine biosynthesis are absent in Chlamydomonas (Hanikenne et al. 2005). In terms of intracellular Cu distribution and storage, the abundant CSDs were suggested to serve as a buffer for cellular Cu levels in yeast and plants since they can sequester Cu and seem to be present in strong excess of the enzymatic activity required to scavenge superoxide species (Culotta et al. 1997; Corson et al. 1998; Cohu et al. 2009). CuZnSODs, however, are missing entirely in Chlamydomonas (Asada et al. 1977; Sakurai et al. 1993; Merchant et al. 2007). Another reason for why we did not observe Cu accumulation under Zn-deficient growth conditions could be that the underlying mechanism in Chlamydomonas depends on CrCRR1-specific protein domains that are not functionally present in AtSPL7. For example, CRR1 contains a pronouncedly cysteine(cys)-rich motif within its C-terminal region which is not as evident in SPL7 (Kropat et al. 2005; Yamasaki et al. 2009). This Cys-rich motif has been associated with CRR1-mediated responses to nickel (Ni) and hypoxia (Sommer et al. 2010; Blaby-Haas et al. 2016). In accordance with a possible lack of such a functional Cys-rich region in SPL7, the response to Ni of Arabidopsis contrasts that of Chlamydomonas. Whereas Ni induced the CRR1-dependent Cu deficiency response in Chlamydomonas even under Cu-sufficient growth conditions, Ni mimicked Cu and suppressed the Cu deficiency response even under low Cu in *A. thaliana* (Kropat et al. 2005; Yamasaki et al. 2009). The Cys-rich motif of CRR1 was additionally implicated in Zn homeostasis because Chlamydomonas cells exhibited constitutive overaccumulation of Zn and transcriptional upregulation of several zinc transporter genes when the Cys-rich domain was removed from the CRR1 protein (Sommer et al. 2010). However, it is presently unknown whether this motif plays a role in CRR1-dependent Cu accumulation under Zn deficiency.

**Figure 5:**
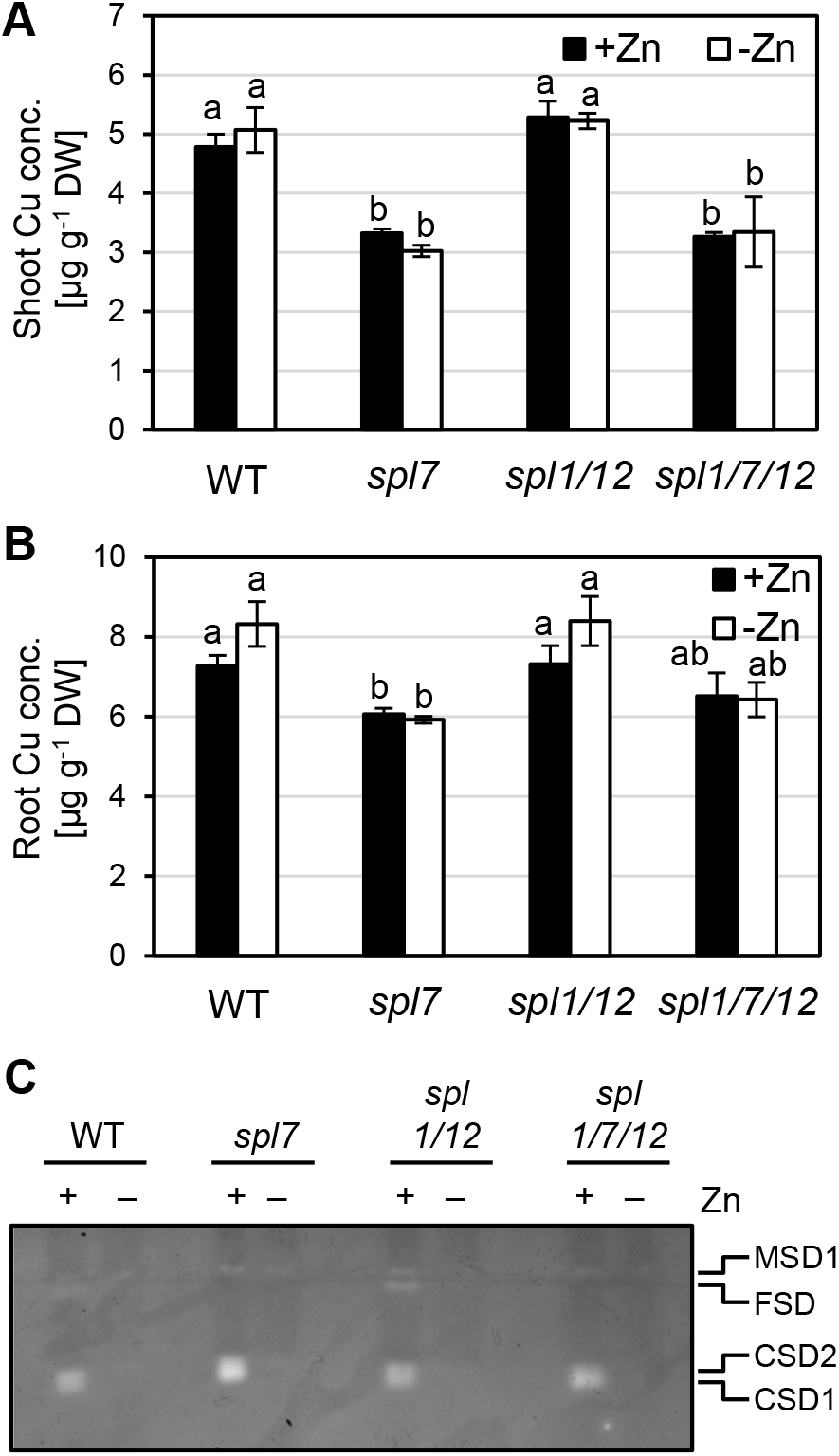
Tissue Cu levels and SOD activities in wild-type, *spl7* single, *spl1 spl12* double and *spl1 spl7 spl12* triple mutant Arabidopsis seedlings cultivated in Zn-deficient and -sufficient media. (A, B) Cu concentrations in shoots (A) and roots (B) of 21-day old seedlings grown on Zn-sufficient (1 μM ZnSO_4_) or Zn-deficient (0 μM ZnSO_4_) agar-solidified media in vertically-oriented plastic petri dishes in short days (11 h). Bars represent arithmetic means ± SD (*n* = 3 replicate plates, each with 20 seedlings. Different characters denote statistically significant differences (*P* < 0.05) between means based on ANOVA (Tukey’s HSD). (C) In-gel detection of superoxide dismutase activities in total protein extracts of seedlings grown as described for (A). Total protein (20 μg per lane) was separated in a 12% (v/v) native polyacrylamide gel. CSD1, CSD2: Cu/Zn Superoxide Dismutases, FSD: Fe Superoxide Dismutase 1-3, MSD1: Mn Superoxide Dismutase 1. Data are from one experiment representative of two to three independent experiments.

We found that *MIR398b* levels were strongly downregulated under Zn-deficient growth conditions, indicating that miR398b is not involved in the downregulation of CSD activities and *CSD2* transcript levels under these conditions (Fig. 6). By contrast, under Cu deficiency a strong SPL7-dependent increase in *MIR398b* and *MIR398c* levels results in reduced levels of *CSD2* transcript levels, a miR398 target, and lowered CSD activity (Shikanai et al. 2003; Yamasaki et al. 2009; Bernal et al. 2012; see Fig. 1). Different from the observations reported here, miR398 was implicated in the decrease of *CSD* transcript levels in roots of Zn-deficient *Sorghum bicolor* (Li et al. 2013). The known isoforms *MIR398a* and *MIR398c* of Arabidopsis were not quantified here and might contribute to the reduction in *CSD* transcript levels under Zn deficiency.

**Figure 6:**
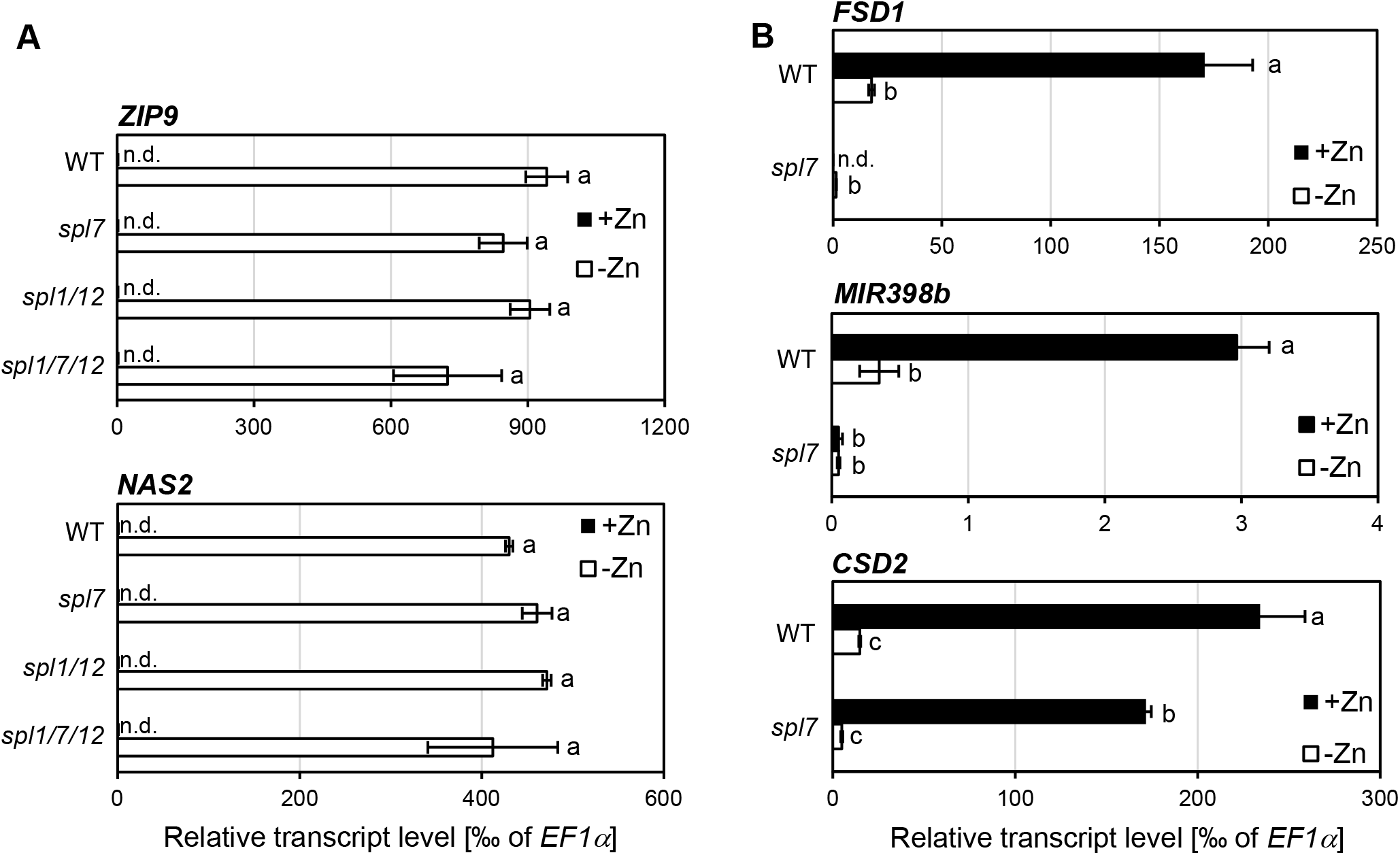
Abundance of marker transcripts in wild-type, *spl7* single, *spl1 spl12* double and *spl1 spl7 spl12* triple mutant Arabidopsis seedlings cultivated in Zn-deficient and -sufficient media. (A, B) Relative transcript abundance, determined by RT-qPCR, of the Zn deficiency markers *ZIP9* and *NAS2* (A) and the Cu deficiency markers *FSD1, CSD2* and *MIR398b* (primary transcript) (B) in 21-day old seedlings grown on Zn-sufficient (1 μM ZnSO_4_) or Zn-deficient (0 μM ZnSO_4_) agar-solidified media in vertically-oriented plastic petri dishes in short days (11 h). Bars represent arithmetic means ± SD (*n* = 3 technical replicates, i.e. independent PCR machine runs, each with three replicate wells per run and transcript). Data shown are transcript levels relative to *EF1α* as a constitutively expressed control gene, from one experiment representative of two independent experiments.

Although we did not observe any connections between Cu and Zn homeostasis in Arabidopsis resembling the molecular manifestations and regulatory mechanisms reported in Chlamydomonas, some crosstalk between the homeostasis of Cu and Zn is known to occur in Arabidopsis (Wintz et al. 2003). In addition, Zn^2+^ and Cu^2+^ ions have similar chemical properties. Metal binding of proteins is predominantly driven by the position of a metal cation in the Irving-Williams series, which predicts that Cu^2+^ can displace Zn^2+^ (Fraústo da Silva and Williams 2001; Waldron et al. 2009; Festa and Thiele 2011), although the type of donor ligands, the properties of the metal-binding pocket or differing preferences of metals for coordination geometries can impart some specificity. These competitive interactions between Zn and Cu could explain some small effects we observed here, possibly including also generally decreased rosette Zn concentrations in *spl7* and *spl1 spl7 spl12* compared to WT (see Fig. S3B). It is important to note that rosette Zn levels remained above critical deficiency Zn concentrations of approximately 20 μg g^−1^ dry biomass in all genotypes (Marschner and Marschner 2012). In conclusion, our results emphasize some differences between metal homeostasis of Chlamydomonas and Arabidopsis.

## Supporting information

Supplemental_information_all

## Acknowledgements

We thank Petra Düchting (Ruhr University Bochum, Germany) for multi-element analysis and Dr. Peter Huijser for discussions and for providing the seeds of the *spl1 spl12* and *spl1 spl7 spl12* mutants. This work was funded by the Deutsche Forschungsgemeinschaft (Kr1967/15-1) and Ruhr University Bochum, Germany.

## Author contributions

AS, MB and UK designed the research; LB, AS and JQ performed research; AS, LB and UK analyzed data; and AS and UK wrote and edited the manuscript with input from the other authors.

## Supplemental Data

Figure S1: Molecular characterization of *spl1 spl12* and *spl1 spl7 spl12* mutants.

Figure S2: Biomass and Cu accumulation of roots of wild-type, *spl7, spl1 spl12* and *spl1 spl7 spl12* mutant seedlings cultivated on Cu-deficient and -sufficient media.

Figure S3: Rosette Cu and Zn accumulation in reproductive-stage wild-type, *spl7, spl1 spl12* and *spl1 spl7 spl12* cultivated on Cu-deficient and -sufficient soil.

Figure S4: Root biomass of wild-type, *spl7, spl1 spl12* and *spl1 spl7 spl12* mutant seedlings cultivated on Zn-deficient and -sufficient media.

Table S1: Oligonucleotides used in this study

Table S2: AGI locus identifiers of genes mentioned in this article

