## Supplemental_information_all for "Do Arabidopsis *Squamosa promoter binding Protein-Like* genes act together in plant acclimation to copper or zinc deficiency?"

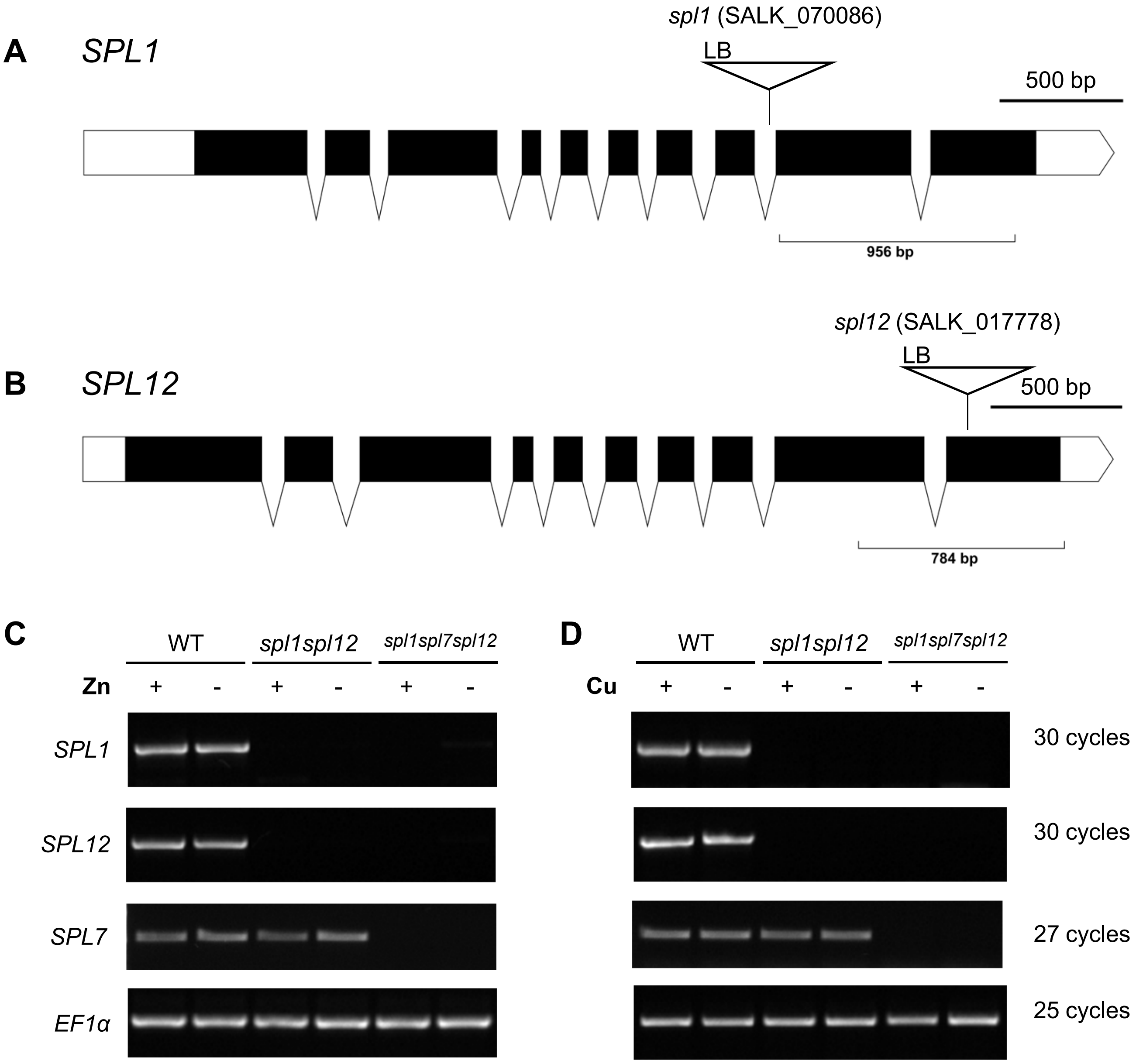


**Figure S1: Molecular characterization of *spl1 spl12* and *spl1 spl7 spl12* mutants.** (A,B) Schematic representation of the *SPL1* (A) and *SPL12* (B) genomic loci. Black boxes represent exons. The positions of the T-DNA insertions marked by open triangles and their orientations given through the positions of left border sequences (LB). The localizations and sizes of the fragments that were amplified in RT-PCRs (see C, D) are marked below (thin black lines). The representation of the genomic loci was generated using the Exon-Intron Graphic Maker (Bhatla 2012). (C, D) RT-PCR analysis of *SPL1*, *SPL7* (507 bp) and *SPL12* transcript levels in wild-type, *spl1 spl12* and *spl1 spl7 spl12* mutant seedlings. Total RNA was extracted from 21-day old seedlings grown either on Zn-sufficient (+, 1 µM $ZnSO_4$) and Zn-deficient (-, 0 µM $ZnSO_4$) agar-solidified media (C) or on Cu-sufficient (+, 0.5 µM $CuSO_4$) and Cu-deficient (-, 0 µM $CuSO_4$) agar-solidified media (D) in short days (11 h). The *EF1α* transcript (fragment of 476 bp) served as a positive control. Note that the reduction of *SPL7* transcript in the *spl7*-2 mutant was confirmed earlier (Bernal *et al.*, 2012). Primer sequences are listed in Supplemental Table 1.

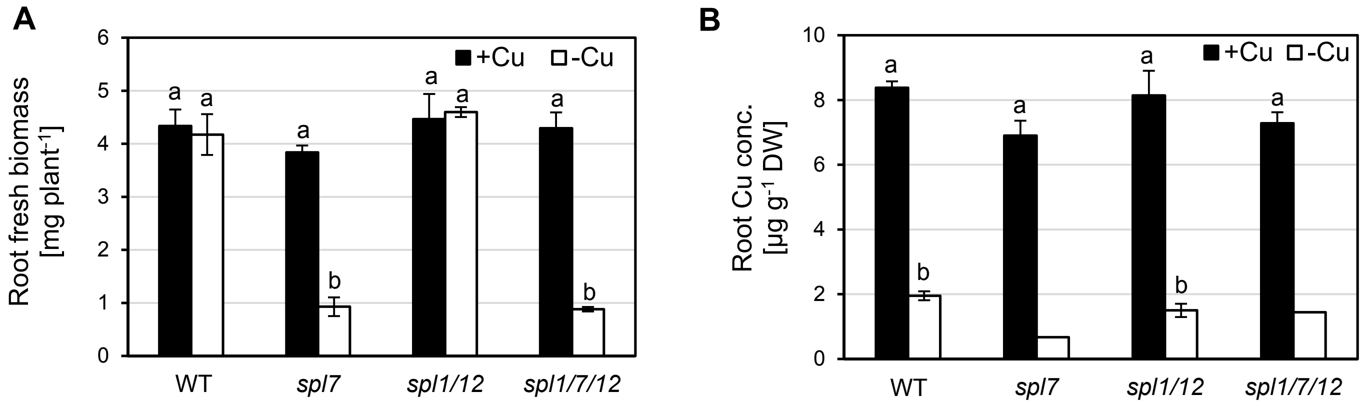


**Figure S2: Biomass and Cu accumulation of roots of wild-type, *spl7*, *spl1 spl12* and *spl1 spl7 spl12* mutant seedlings cultivated on Cu-deficient and -sufficient media.** Root biomass (A) and root Cu concentrations (B) of 21-day-old seedlings grown on Cu-sufficient (0.5 µM $CuSO_4$) or Cu-deficient (0 µM $CuSO_4$) agar-solidified media in vertically-oriented glass plates in short days (11 h). Bars represent arithmetic means ± SD ($n$ = 3 replicate plates, each with 20 seedlings). In (B), no SD is shown for *spl7* and *spl1 spl7 spl12* under -Cu in because the dry root biomass was so low that all three replicate plates were pooled to obtain one value. Different characters denote statistically significant differences ($P < 0.05$) between means based on ANOVA (Tukey's HSD). Data are from one experiment representative of three independent experiments.

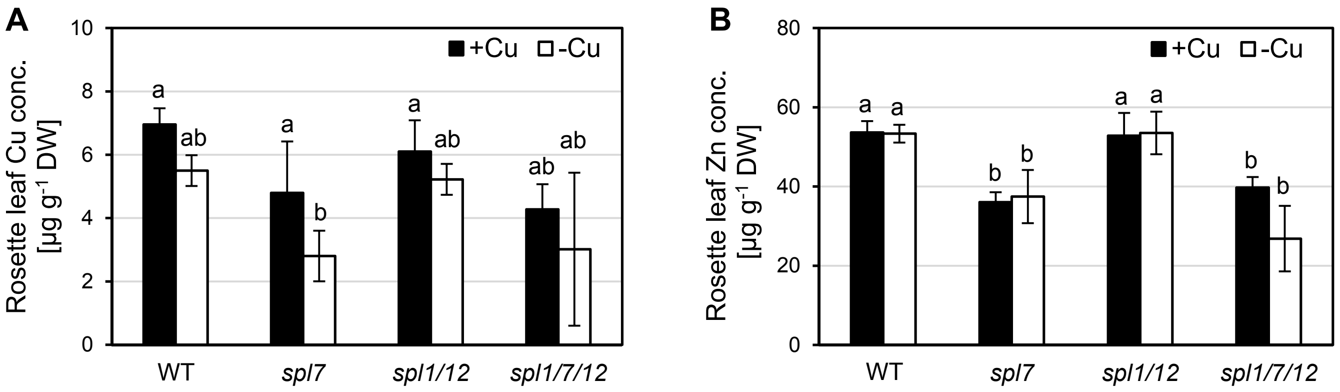


**Figure S3: Rosette Cu and Zn accumulation in reproductive-stage wild-type, *spl7*, *spl1 spl12* and *spl1 spl7 spl12* cultivated on Cu-deficient and -sufficient soil.**

Cu concentration (A) and Zn concentration (B) in rosette leaves of 40-day old plants watered with equal amounts of either tap water or freshly prepared 2 mM $CuSO_4$ in tap water and grown in long days (16 h). Bars represent arithmetic means ± SD (*n* = 6 individual plants per genotype and treatment). Different characters denote statistically significant differences ($P < 0.05$) between means based on ANOVA (Tukey's HSD). Data are from one experiment representative of two independent experiments.

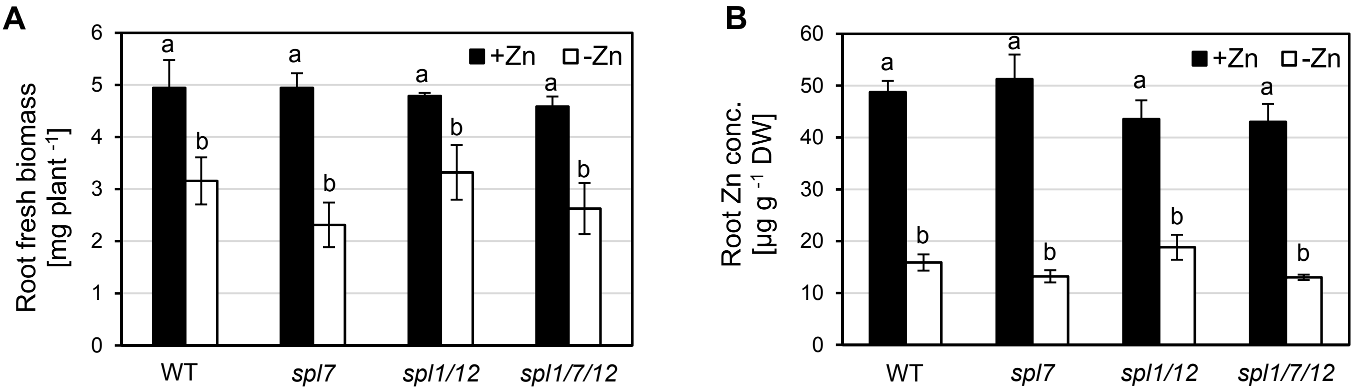


**Figure S4: Root biomass of wild-type, *spl7*, *spl1 spl12* and *spl1 spl7 spl12* mutant seedlings cultivated on Zn-deficient and -sufficient media.**

(A) Root biomass of 21-day-old seedlings on Zn-sufficient (1 µM $ZnSO_4$) or Zn-deficient (0 µM $ZnSO_4$) agar-solidified media grown in vertically-oriented plastic petri dishes in short days (11 h). Bars show arithmetic means ± SD ($n$ = 3 replicate plates, each with 20 seedlings). Different characters denote statistically significant differences ($P$ < 0.05) between means based on ANOVA (Tukey's HSD). Data are from one experiment representative of three independent experiments.

### Table S1: Oligonucleotides used in this study

| Oligo name | Oligo sequence (5' → 3') | Reference |
|---|---|---|
| **Oligos used for genotyping & RT-PCR** | | |
| At_spl1_geno_f | CCCCTCTTTGACAATACTCGTC | this work |
| At_spl1_geno_r | GGATGTATGAGAAGTGACCGCG | this work |
| At_spl12_geno_f | GGCACTGTTGATCCATCTCCTGATGCTGCG | this work |
| At_spl12_geno_r | GGTATAGGGAAGTTTTACTAGCTTGTTCC | this work |
| LB_T-DNA | AACGTCCGCAATGTGTTATTAAGTTGTC | Woody et al. 2007 |
| AtSPL1_RT_f | GACTCAGTAGCAGCTTCTTCC | Schwarz 2006 |
| AtSPL1_RT_r | CACACAGACGCAAACCGCAGC | Schwarz 2006 |
| AtSPL7_RT_f | CTATTCTGTTGTACCTGCACCG | this work |
| AtSPL7_RT_r | CAGTGAACAAGACTGTCTGGC | this work |
| AtSPL12_RT_f | GGTATAGGGAAGTTTTACTAGCTTGTTCC | Schwarz 2006 |
| AtSPL12_RT_r | GGCACTGTTGATCCATCTCCTGATGCTGCG | Schwarz 2006 |
| AtEF1α_RT_f | TAAGGATGGTCAGACCCGTGA | Sinclair et al. 2018 |
| AtEF1α_RT_r | CAGACTCGTGGTGCATCTCAAC | Sinclair et al. 2018 |
| **Oligos used for qRT-PCR** | | |
| AtEF1α_qRT_f | TGAGCACGCTCTTCTTG | Czechowski et al. 2005 |
| AtEF1α_qRT_r | GGTGGTGGCATCCATCT | Czechowski et al. 2005 |
| AtFSD1_qRT_f | TCGGCTCTTTCCATTGCTT | Bernal et al. 2012 |
| AtFSD1_qRT_r | TGGTCTTCGGTTCTGGAAGTCA | Bernal et al. 2012 |
| AtNAS2_qRT_f | CTGACGACGTGGTTAATTCGG | Talke et al. 2006 |
| AtNAS2_qRT_r | TGCCTCGAGCTCCATTTGA | Talke et al. 2006 |
| AtpriMIR398b_qRT_f | CACGAGTAATCAACGGCTGTAATG | Bernal et al. 2012 |
| AtpriMIR398b_qRT_r | TGAGTAAAAGCCAGCCTTGATAAAAG | Bernal et al. 2012 |
| AtZIP9_qRT_f | CCATCACTACTCCGATCGGTGT | Talke et al. 2006 |
| AtZIP9_qRT_r | CACCAATGCTGCAACGCTATAA | Talke et al. 2006 |

**Table S2: AGI locus identifiers of genes mentioned in this article**

| **Gene abbreviation** | **AGI locus identifier** |
|---|---|
| *CITF1* | AT1G71200 |
| *CSD1* | AT1G08830 |
| *CSD2* | AT2G28190 |
| *FSD1* | AT4G25100 |
| *HMA5* | AT1G63440 |
| *IRT1* | AT4G19690 |
| *MIR398b* | AT5G14545 |
| *MSD1* | AT3G10920 |
| *NAS2* | AT5G56080 |
| *PC* | AT1G76100, AT1G20340 |
| *SPL1* | AT2G47070 |
| *SPL2* | AT5G43270 |
| *SPL7* | AT5G18830 |
| *SPL8* | AT1G02065 |
| *SPL10* | AT1G27370 |
| *SPL11* | AT1G27360 |
| *SPL12* | AT3G60030 |
| *SPL14* | AT1G20980 |
| *SPL16* | AT1G76580 |
| *ZIP9* | AT4G33020 |
